# The Trade-Off Between Torque and Power with Speed: A Study of Shoulder Performance During an Isokinetic and Multiplanar Task

**DOI:** 10.1101/2024.11.06.616796

**Authors:** Kayla D.L Lee, Erin C.S Lee, Michael J. Rainbow

## Abstract

The human shoulder likely evolved under selective pressures favouring diverse tasks that require high mobility, speed, and torque. For example, humans are uniquely adept at high-speed and accurate throwing. Prior work has aimed to quantify the kinematics and kinetics of upper limb movements in isometric or uniplanar motions. However, we still do not fully understand the trade-offs of shoulder torque and power with angular velocity during functional tasks that are reflective of demands that may be relevant to the shoulder’s evolution. We developed a novel approach for upper limb 3D inverse dynamic calculations by integrating motion capture with an instrumented cable machine. Twenty-five participants performed a crossbody, isokinetic upper limb motion at various cable speeds in a rigid and free torso condition (self-imposed). Shoulder torque decreased significantly (p < 0.05) with increasing angular velocity in 19 and 16 participants for the constrained and unconstrained conditions, respectively. Shoulder power increased significantly (p < 0.05) with angular velocity for 6 and 11 participants for constrained and unconstrained, respectively. T-tests revealed no statistical difference between the torso conditions for torque and power against angular velocity. Our findings suggest that despite having a trade-off in torque and velocity, the shoulder may be tuned to produce power over a wide range of velocities independent of energy transfer from the lower extremities.

## Introduction

The human shoulder is an intricate joint-complex comprised of four articulations. The shoulder also consists of 19 muscles that span the joint, facilitating a range of motion that covers approximately 65% of a sphere which allows the hand to be positioned almost anywhere around the body (Larson, 2009; Veeger and Van Der Helm, 2007). These features make the shoulder the most mobile and fastest joint complex in the human body (Pappas et al., 1985; Roach et al., 2013). Potential selective pressures (such as tool-making, digging, climbing, combat, and load-carrying) throughout our evolution may have shaped the human shoulder to be capable of diverse tasks requiring wide-ranging combinations of mobility, speed, and strength (Atwater, 1979; Larson, 2009; Veeger and Van Der Helm, 2007). In particular, hunting prey and defending kills throughout our hominin lineage presumably required frequent throwing of hand-cast projectiles based on archaeological evidence of primitive weapons (Shea, 2006). Our unique ability to overhand throw projectiles for distance, speed, and accuracy requires high torque and velocity delivered over a large range of motion (Isaac, 1987; Lombardo and Deaner, 2018; Roach and Lieberman, 2014).

During high-speed throwing, the shoulder is an integral component of the whole-body kinetic chain. Beginning at the foot, carefully sequenced activation and rapid contraction of muscles generate torques at each joint with specific timing. As a segment accelerates and decelerates, the subsequent segment obtains the greatest speed of the segment preceding it, then accelerates to an even faster speed. This pattern progresses from the planted foot, through the pelvis and torso, then the shoulder, elbow, and wrist of the throwing arm, resulting in the highest velocity at the hand during the time of release (Atwater, 1979; Hirashima et al., 2002; Pappas et al., 1985; Putnam, 1993; Roach and Lieberman, 2014). The shoulder’s own torque generation over its large range of motion supplements the velocity from the preceding joints, and effectively transmits energy from the large lower body and trunk segments to the end of the hand (Ben Kibler, 1998). Shoulder torque, angular velocity, and power have been studied during pitching (Lin et al., 2003; Roach and Lieberman, 2014; Roach et al., 2013); however, due to variances in the timing of the pitching phases and the overall rapidity of the throw, the torque-velocity and power-velocity relationships of the shoulder are challenging to characterize.

The relationship between torque and angular velocity in musculoskeletal joints is a trade-off reflecting the well-established force-velocity relationship of contracting muscles: as velocity increases, the generated force decreases (Fenn and Marsh, 1935; Tihanyi et al., 1982; Wilkie, 1949; Zaciorskij et al., 2005). The inverse relationship between torque and velocity has been studied for joints with fewer degrees of freedom, such as the knee and elbow (Borges et al., 2003; Harries and Bassey, 1990; Kannus and Beynnon, 1993; Taylor et al., 1991); however, less is known about the shoulder. Studies have quantified the shoulder’s torque and angular velocity relationship in isometric (Baillargeon et al., 2022; Leonardis et al., 2020; Moghadam et al., 2011; Otis et al., 1990) or uniplanar motions (Koski and McGill, 1994; Whitcomb et al., 1995; Wight et al., 2022; Zanca et al., 2011), and results have been similar to those of other joints. However, they do not reflect the shoulder’s ability to perform complex, three-dimensional, and high-speed movements. More functional actions, such as the bench press exercise, have also been studied to evaluate the force and velocity of the upper limb (Cosic et al., 2021; Rahmani et al., 2018); however, the specific contribution of the shoulder and the joint kinetics during the multi-joint movement were not a primary focus.

Mechanical power is calculated as the dot product of force and velocity (P = F · v) in linear cases, such as for muscles, or as the product of torque and angular velocity (P = τ · ω) in rotational cases, such as for joints. Muscle power (linear) and joint power (rotational) are different expressions of mechanical power, useful for demonstrating the rate of mechanical energy and overall work being done. Power production is generally high in animals performing fast movements (Lappin et al., 2006; Patek et al., 2010; Roberts and Scales, 2002; Sutton et al., 2022), including in human throwing (Roach et al., 2013; Wilk et al., 1993). The theoretical power-velocity relationship exhibits power increasing with velocity to a peak at an intermediate contraction velocity, where a combination of force and velocity yields the highest power output. Then, power declines as contraction velocity continues to increase and force production decreases (Zaciorskij et al., 2005). Previous research has suggested that rotation of the torso passively loads the elastic, soft tissue components in the shoulder for energy storage during arm-cocking phase of the throwing sequence. The release of elastic energy produces a large portion of the necessary shoulder power for the rapid internal rotation and anterior acceleration of the humerus during the acceleration phase of the throwing sequence. The same study also estimated that ∼30% of shoulder power is produced by the torso and hip rotators (Roach et al., 2013). Given that the shoulder can generate high torques while also producing the fastest angular velocities in the body; 800 deg s ^-1^ in horizontal adduction (Seminati et al., 2015), 1000 deg s ^-1^ in flexion (Jessop and Pain, 2016; Wagner et al., 1992), and 9000 deg s ^-1^ in internal rotation (Feltner and Dapena, 1986; Pappas et al., 1985), we aim to explore how power and speed may trade off. Since prior studies focusing on the lower limb have only reported values on the ascending limb of the power-velocity curve for joint power during knee extension without reaching tested angular velocities on the descending limb (Aagaard et al., 1994; Lanza et al., 2003; Taylor et al., 1991), we are seeking to test angular velocities in the shoulder where we may be able to view power peaking or declining.

These scientific gaps are partly due to the technical challenges and intricacies of collecting high-quality data for such a task that also allows inverse dynamics to be computed for the upper limb. Inverse dynamics is a branch of mechanics that combines kinematics (motion without considering forces) with kinetics (forces that are caused or produced by motion) and is the primary method for obtaining detailed information about the net forces and torques produced at each joint during movement (Robertson et al., 2004). There is considerable research that uses inverse dynamics for the lower limb, where force plates capture the ground reaction forces at the foot to then calculate the joint forces up the lower limb chain (e.g. Armstrong et al., 2022; Caruntu and Moreno, 2019; Cluff et al., 2008; Franz and Kram, 2014; Ren et al., 2008). This option for the shoulder is not optimal as a significant amount of error is introduced through propagation with each added segment moving up the kinetic chain (Vlietstra, 2014), and the split of forces through the torso to each arm is challenging. An alternative method is to compute inverse dynamics from the hand, as fewer segments are involved. This approach has been done for baseball pitching using the interaction of the hand and ball as the external force (Aguinaldo et al., 2007; Dowling et al., 2020; Feltner and Dapena, 1986; Roach and Lieberman, 2014), exercises using constant and elastic loads (Häberle et al., 2018), and simulated workplace tasks (Mulla et al., 2020). Still, there remains a gap in understanding the kinematics and kinetics of multiplanar upper limb movement across angular velocities across a wide range of speeds.

This study had two objectives; (1) to characterize the torque-velocity and power-velocity relationships for the shoulder during a high-demand and multiplanar movement, and (2) to test how restricting the torso’s rotation while performing a multiplanar shoulder motion would affect the torque and power generation. For the first objective, we anticipated that shoulder torque would decrease with increasing angular velocity, expressed as negative slopes statistically different than zero and reflecting the force-velocity relationship in muscles. For the second objective, we hypothesized that restricting torso rotation would result in shoulder torque decreasing faster with increasing velocity than when the torso is recruited (natural movement). To accomplish our objectives, we computed kinematics and kinetics at the shoulder using an approach that combines markerless and marker-based motion capture with an actuated cable system.

## Materials and Methods

### Participants

Twenty-five healthy participants (13F/12M, age 22 ± 3 years; body weight 69 ± 11 kg; height 172 ± 9 cm) were recruited for this study. Eligibility criteria included no upper extremity injury in the last year and no neuromuscular disorders. Ethics approval was granted by the Queen’s University General Research Ethics Review Board. All participants were informed of the testing protocol and provided written informed consent to participate in the study and for their data to be used for research.

### Experimental procedure

The participants performed a crossbody arm motion with their right arm against a tensile load supplied by an instrumented cable machine (1080 Quantum, Sweden) typically used in high-performance training centers. We gave participants verbal instructions and visual cues to perform the motion. Beginning with their right arm reaching overhead to the hanging cable to their right (Figure 1A), they were instructed to pull the cable across their body while maintaining a straight arm to end at their left hip (Figure 1B). This movement pattern was chosen as it exhibited a large range of upper limb motion and moved through all planes of the body. The machine allowed us to collect data in an isokinetic setting. Using the isokinetic setting, the 1080 supplied the required force to maintain a prescribed constant top-speed. The 1080 machine is capable of concentric speeds ranging from 0.05 m s ^-1^ to 8 m s ^-1^. We co-calibrated 8 markerless (Miqus) and 8 marker-based (Oqus) cameras to collect simultaneous 3D kinematics of the participant (Theia3D, Canada) and marker tracking of the cord (Qualysis, Sweden) at 120 Hz.

**Figure 1:**
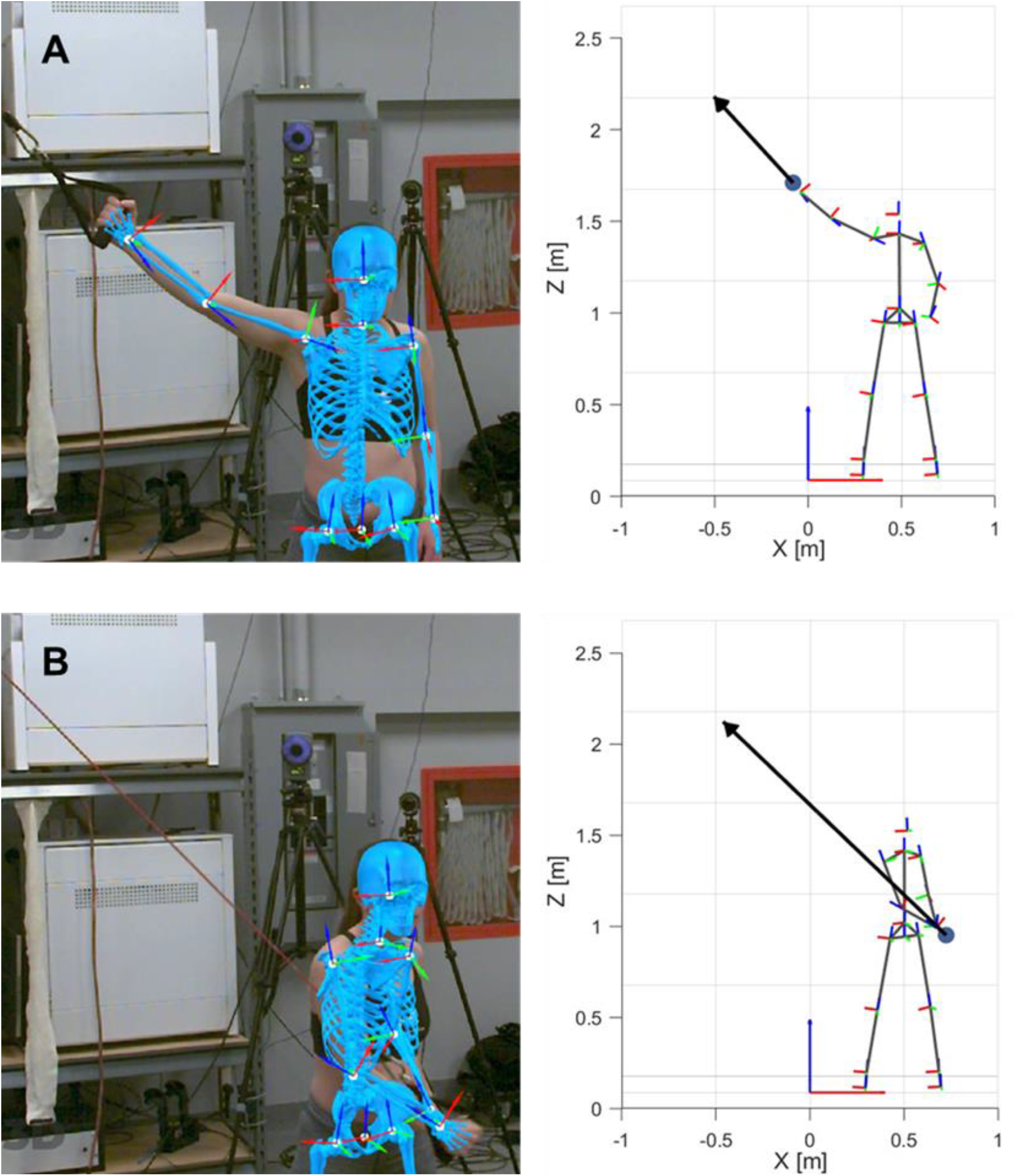
Start and end frames of the task during a constrained trial. The 3D skeleton was produced in Theia3D (left). The link-segment model was produced in MATLAB based on the kinematics from the Theia3D output and the force vector was generated from the motion capture and cable machine outputs (right). The large sphere at the base of the force vector represents the middle of the hand.

To test the hypothesis that shoulder torque decreases faster when the lower body and torso are not engaged, the participants performed the task in two self-imposed torso conditions; constrained and unconstrained (Figure 2). In the constrained condition, we instructed participants to keep the torso relatively rigid throughout the movement. For the unconstrained condition, they were allowed to rotate their torso while limiting lower leg use.

**Figure 2:**
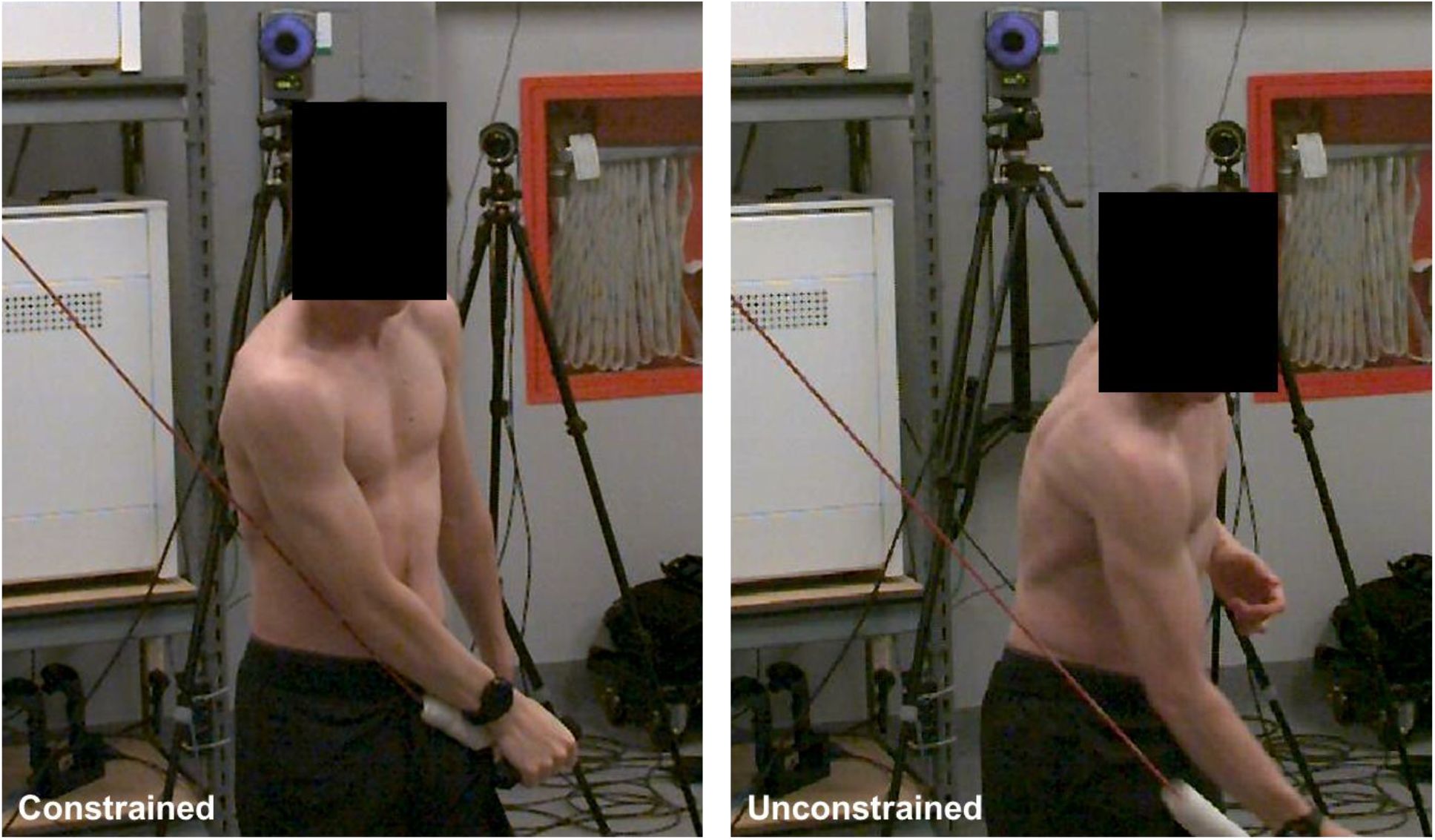
The constrained and unconstrained torso conditions were self-imposed during the task. These frames show the end range of the task to highlight the difference in axial rotation of the trunk.

The main data collection was preceded by a guided warm-up routine consisting of 6 exercises to activate the shoulder, chest, and upper back muscles. We then tested the participants’ maximum cable speed. This test consisted of a single repetition of the task as fast as possible with the cable load set to 1 kg (lowest) and the maximum cable speed set to the system’s maximum (8 m s ^-1^). Using the maximum speed result, we randomized test speeds ranging from 1.5 m s ^-1^ to the participant’s peak cable speed plus 1 m s ^-1^ – rounded to the nearest 0.5 increment for the main collection.

Using the maximum speed result, we randomized test speeds ranging from 1.5 m s ^-1^ to the participant’s peak cable speed plus 1 m s ^-1^ with increments of 0.5 m s ^-1^ for the primary collection. The primary data collection consisted of testing each of the cable speeds from the randomized order. The order for the torso condition was also randomized among participants. With each test speed, we informed the participants of the cable speed to be tested. They were permitted 1-3 practice repetitions to gauge the cable speed and could opt to skip the practice if chosen. For each trial, the participants performed 3 repetitions of the task while we verbally encouraged them to apply maximal effort throughout the entire range of motion. Between each trial, the participants rested for a minimum of 30 seconds. Once all test speeds were completed for one torso condition, the same procedure was repeated for the 2^nd^ condition with a new order of randomized speeds.

To assess whether fatigue or learning effects were present in post hoc analysis, each collection began and ended with a maximum force test in addition to the maximum speed test. During the maximum force test, the maximum cable speed was set to 0.05 m s ^-1^ (the system’s minimum) and participants were asked to perform the motion with as much force as possible. The participants were allowed to rest for a self-selected time after the maximum tests, before proceeding with the main data collection.

### Data processing and calculation of biomechanical metrics

To obtain coupled shoulder kinematics and kinetics, we integrated the marker-based and markerless motion capture data with the force output from the 1080 machine. We conducted residual analysis to determine the cutoff frequency with iterations between 5 and 60 Hz (Nyquist frequency) for data filtering. Based on this analysis, the kinematic data was processed and filtered at a cutoff frequency of 8 Hz in Theia3D. We time-synchronized the 1080’s and motion capture by using the 1080’s cable position output and the reflective markers placed along the cable for the cameras. We calculated the force vector applied to the hand from the force magnitude measured by the 1080 machine and the line of action made by the cable markers when pulled taught. In MATLAB, we performed 3D inverse dynamics to compute shoulder torque, angular velocity, power, and joint angles. We used Dempster’s body segment distribution ratios to approximate the segment masses of the upper arm, lower arm, and hand as 0.6%, 1.6%, and 2.8% of body weight, respectively (Dempster, 1955). Since we were concerned with the torque responsible for causing the motion, we used the component of shoulder torque aligned with the instantaneous angular velocity vector. We parsed each of the three reps in each trial and narrowed frames of interest to only the concentric phase of the movement. To do this, the start of each rep was indicated by the acceleration of the hand exceeding a specified threshold. The end event was detected using the hand position; due to the nature of the task, the end of the concentric phase coincided with the instant the hand was at its lowest vertical and left-most horizontal position. The detected events agreed with the start and end of the cable machine’s signal. We selected values for torque and angular velocity that corresponded to the instant of peak power to assess how our metrics varied across trials (Figure 3). Typically, torque and angular velocity are extracted at a specific joint angle across all trials; however, we opted for the instant of peak power to allow variation and find the point where the angular velocity and torque were both high, regardless of where it occurred along the motion path. We used the rep with the highest peak power value for each trial.

**Figure 3:**
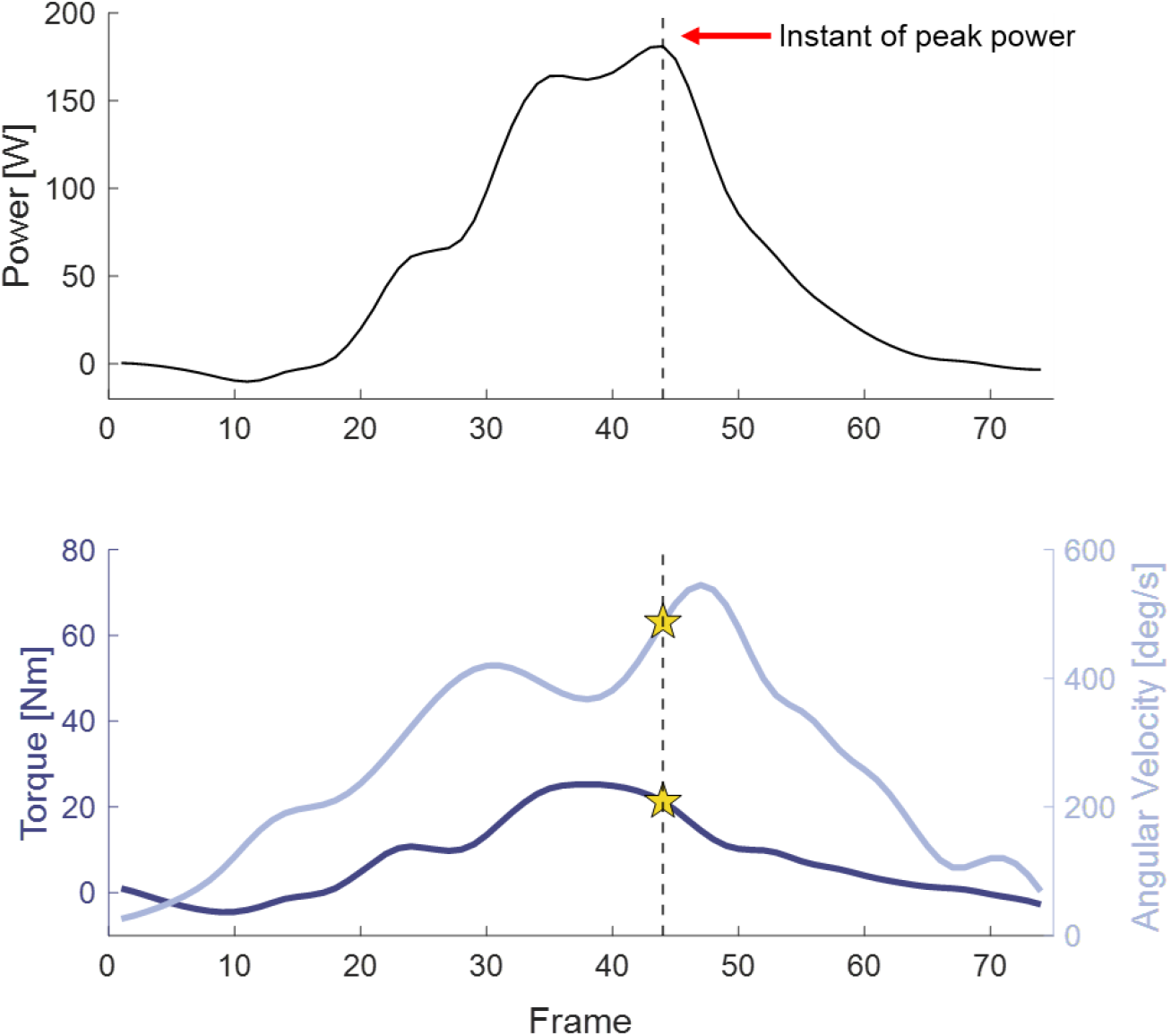
Values for shoulder torque and angular velocity were determined based on the instant of peak power for each trial.

### Analysis and Statistics

We applied linear regression to assess the relationships between the shoulder torque and power with angular velocity. Paired t-tests were used to determine the within-participant differences between conditions for torso rotation, torque-velocity slopes, power-velocity slopes, and the maximal effort tests. In addition, when analyzing the data at the slowest and fastest angular velocities, we used the experimental values rather than the predicted values from the linear regressions.

## Results

### Torso Conditions

Although the torso conditions were self-imposed, rotation about each axis was significantly reduced (p < 0.001) between the unconstrained to constrained conditions; from 96° (SD = 19°) to 51° (SD = 14°) in the long axis of the torso (transverse plane), from 28° (SD = 9°) to 17° (SD = 9°) in lateral flexion (frontal plane), and from 31° (SD = 11°) to 16° (SD = 6°) in flexion/extension (sagittal plane) (Figure 4).

**Figure 4:**
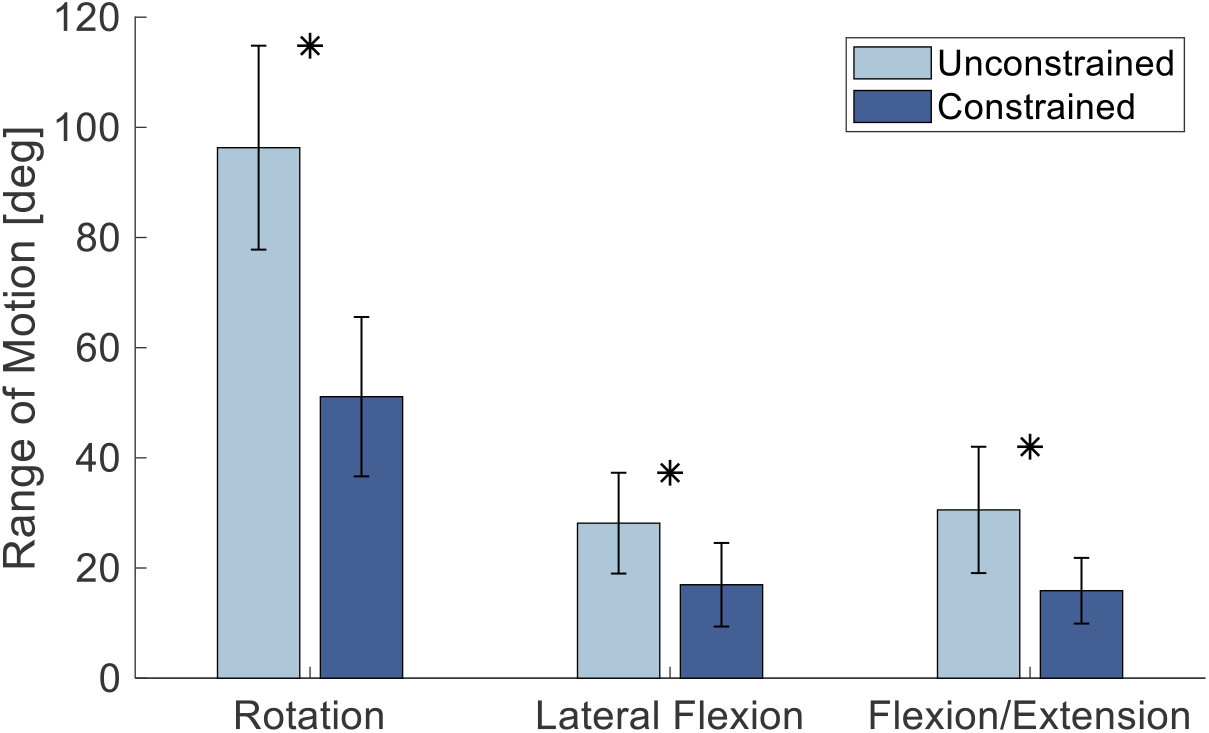
Range of motion for self-imposed torso conditions. The asterisk denotes statistically different pairs; p<0.01e-6 for all axes of rotation.

### Shoulder Torque and Angular Velocity

Across participants, the mean slowest angular velocity for the constrained condition was 234 deg s ^-1^ and had a range of 143 – 469 deg s ^-1^, while the fastest angular velocity averaged 574 deg s ^-1^ with a range of 376 – 784 deg s ^-1^ (Figure 5A). For the unconstrained condition, the mean slowest angular velocity across all participants was 223 deg s ^-1^ with a range of 148 – 343 deg s ^-1^, and the mean fastest angular velocity was 506 deg s ^-1^ with a range of 391 – 671 deg s ^-1^ (Figure 5B).

**Figure 5:**
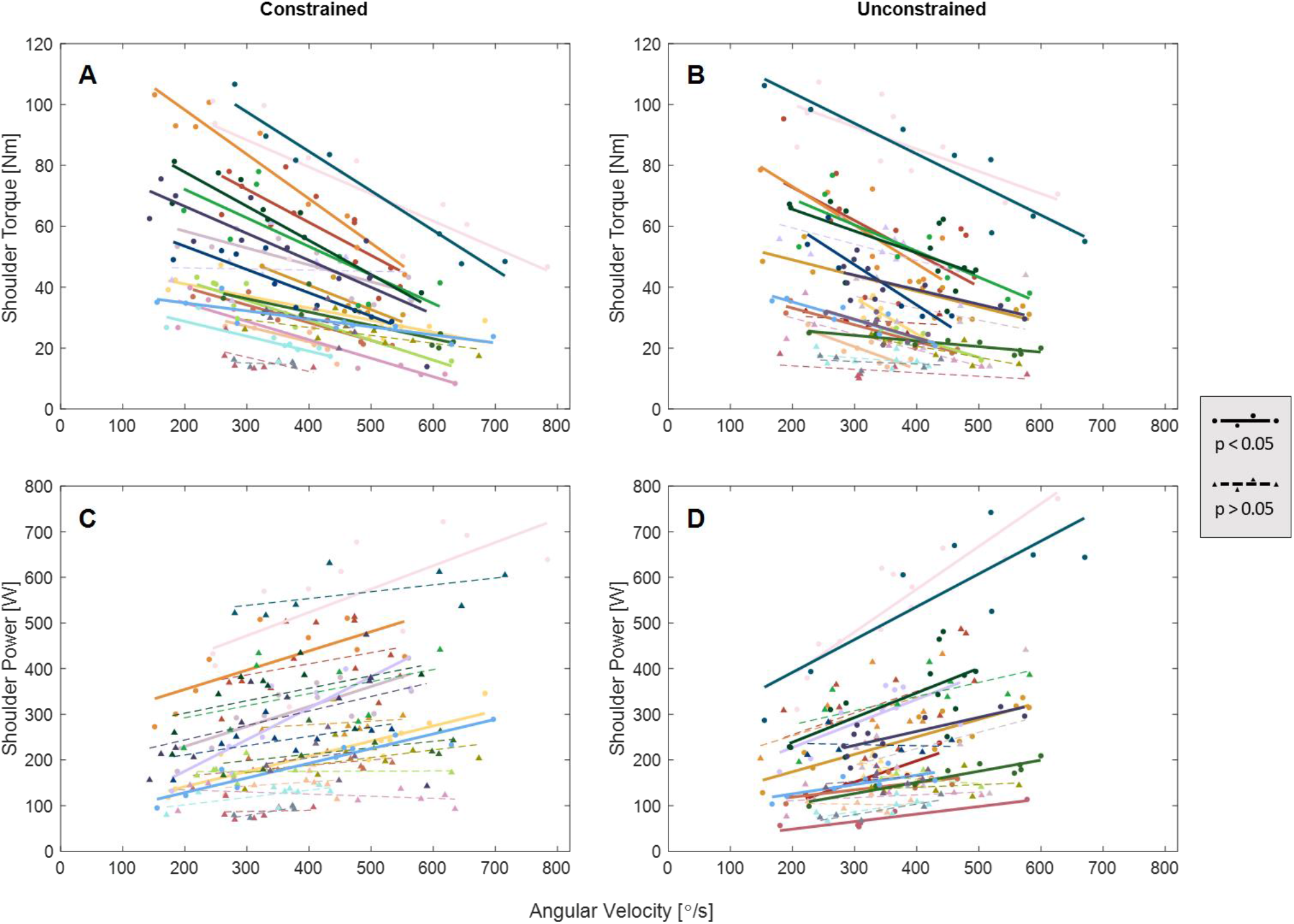
Shoulder torque and power as a function of angular velocity for the constrained and unconstrained conditions.

All participants decreased their torque as angular velocity increased regardless of the condition (Figure 5A, B). Among 24 participants (1 exclusion due to large gaps in data), there were 19 and 16 individuals with significant (p < 0.05) relationships for the constrained and unconstrained conditions, respectively. Of these statistically significant torque-velocity slopes, both conditions averaged -0.08 Nm deg ^-1^ s ^-1^ (R^2^ = 0.7 and R^2^ = 0.6 for constrained and unconstrained, respectively).

### Shoulder Power

Of the 24 participants, power increased as angular velocity increased in 22 and 20 participants for constrained and unconstrained, respectively (Figure 6). There were 6 and 11 significant (p < 0.05) power-velocity relationships with average slopes of 0.5 W deg ^-1^ s ^-1^ (SD = 0.04 W deg ^-1^ s ^-1^) and 0.4 W deg ^-1^ s ^-1^ (SD = 0.04 W deg ^-1^ s ^-1^) (both R^2^ = 0.6) for constrained and unconstrained, respectively (solid lines) (Figure 5C-D, Figure 6).

**Figure 6:**
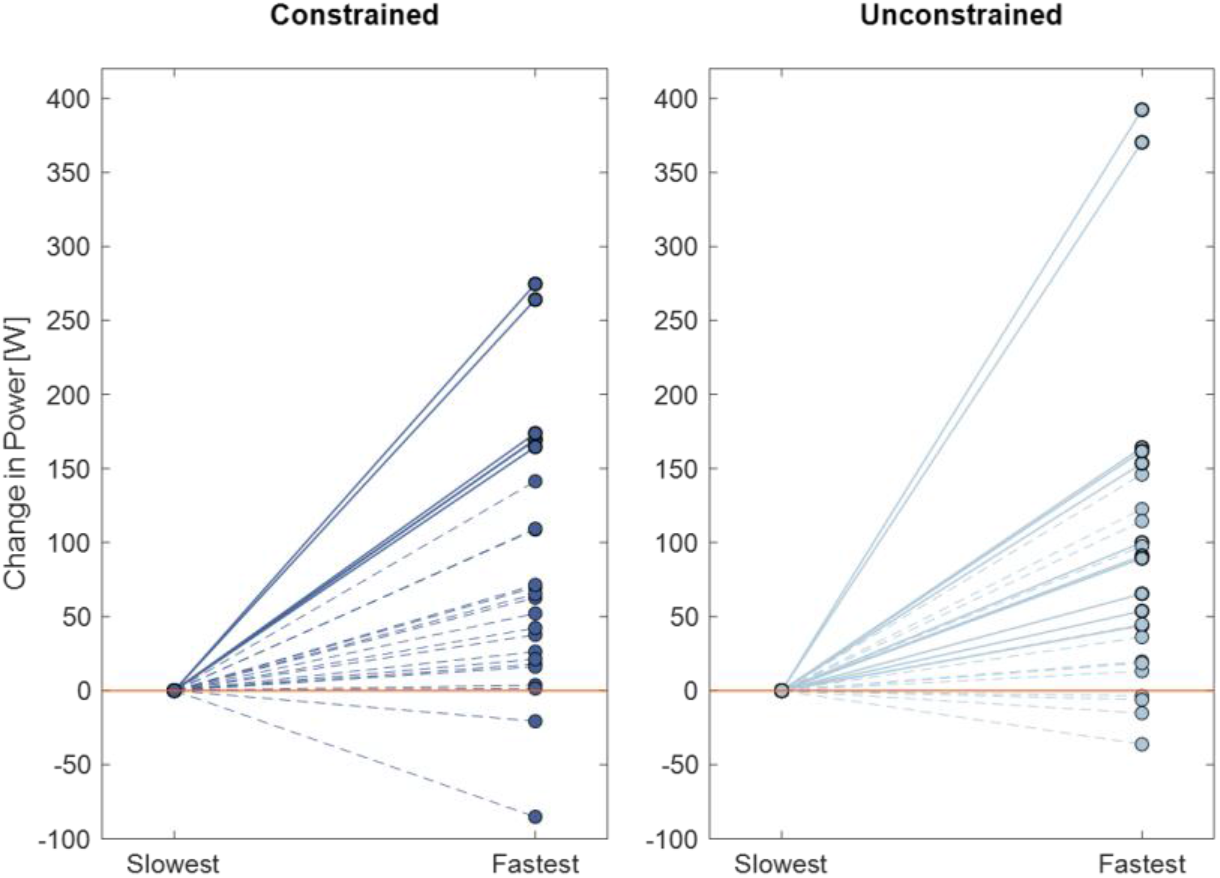
Change in power that occurs at the slowest and fastest angular velocity for each participant. Solid lines indicate participants with a statistically significant power-velocity relationship.

### Pre- and post-collection maximal effort tests

Across participants, the average peak force during the isometric test was 153 N (SD = 47 N) and 162 N (SD = 57 N) for pre- and post-collection, respectively (Figure 7, left). A paired t-test showed no statistical difference (p = 0.29) between the peak forces recorded before and after the data collection. The pre- and post-collection results for the maximum speed tests averaged 5.2 m s ^-1^ (SD = 1.1 m s ^-1^) and 5.7 m s ^-1^ (SD = 1.2 m s ^-1^), respectively (Figure 7, right), where the maximum speed was significantly higher in post-collection than in pre-collection (p < 0.001).

**Figure 7:**
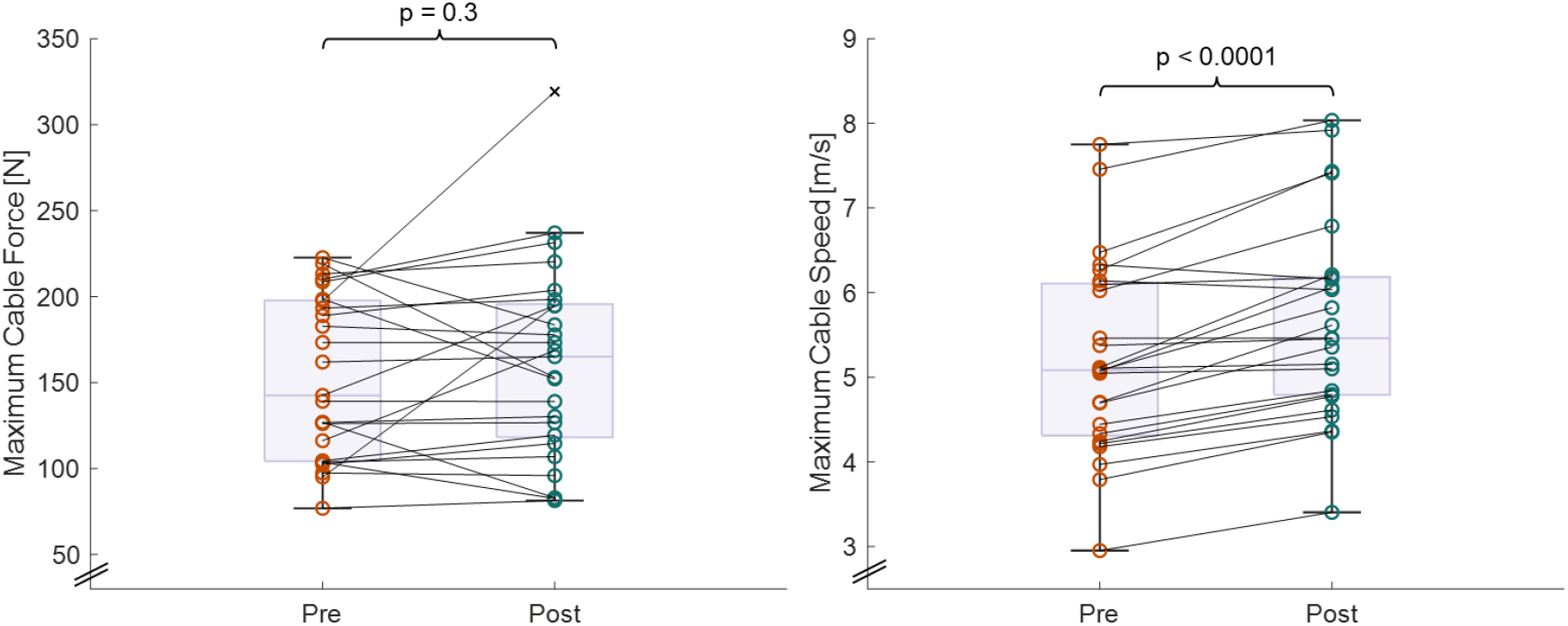
Maximum recorded cable force (left) and cable speed (right) collected before and after the main data collection.

## Discussion

We used a novel approach integrating markerless motion capture and a cable-actuated force measurement system to characterize the torque-velocity and power-velocity relationships of the shoulder during a loaded, dynamic, and multiplanar task. We found that shoulder torque decreased with increasing angular velocity (agreeing with our first objective’s hypothesis) for all participants regardless of the torso condition (contrasting with our second objective’s hypothesis) and exhibited torque decreasing more gradually with angular velocity compared to reported values for the lower limb (Borges et al., 2003; Harries and Bassey, 1990; Pontaga, 2004; Wagner et al., 1992). The power-velocity relationships had no statistical difference between conditions and 22 and 20 participants increased power generation with increasing angular velocity for constrained and unconstrained, respectively. This indicates that absolute maximal power may not have been achieved despite the presence of a torque-velocity trade-off.

### Torque-velocity relationship

We observed that torque decreased as angular velocity increased for all participants. This relationship aligns with the well-researched properties of the force-velocity relationship in muscles and with literature studying multi-joint muscle output during maximal performance tasks such as vertical jumping (Cuk et al., 2014; Zivkovic et al., 2017), squatting (Armstrong et al., 2022; Bosco et al., 1995; Rahmani et al., 2001), cycling (Bozic and Berjan Bacvarevic, 2018), bench pressing (Rahmani et al., 2018; Zivkovic et al., 2017), and wheelchair sprinting (Hintzy et al., 2003). The relationship between joint torque and angular velocity is comparable to the inverse relationship between muscle force and contraction velocity. While existing literature focusing on coupled joint kinematics and kinetics has predominantly focused on the lower limb (Borges et al., 2003; Hahn et al., 2014; Harries and Bassey, 1990; Taylor et al., 1991; Wagner et al., 1992; Yeadon et al., 2006) and the shoulder in uniplanar (Koski and McGill, 1994; Whitcomb et al., 1995; Wight et al., 2022; Zanca et al., 2011) or isometric contexts (Baillargeon et al., 2022; Leonardis et al., 2020; Moghadam et al., 2011; Otis et al., 1990), we observed torque-velocity profiles that generally align with these prior studies during our multiplanar task. The mean slopes of the linear regressions across participants (−0.08 Nm deg ^-1^ s ^-1^ and -0.07 Nm deg ^-1^ s ^-1^ for constrained and unconstrained, respectively) are comparable to the estimated torque-velocity slopes for arm flexion (−0.07 Nm deg ^-1^ s ^-1^) (Hortobagyi and Katch, 1990), shoulder abduction in the coronal (−0.07 Nm deg ^-1^ s ^-1^) and scapular (−0.06 Nm deg ^-1^ s ^-1^) planes (Whitcomb et al., 1995), and shoulder flexion (−0.05 Nm deg ^-1^ s ^-1^) (Martins et al., 2024). These slopes for the upper extremity appear to be less steep than those of the lower extremity which prior studies report being approximately between 0.1 to -0.5 Nm deg ^-1^ s ^-1^ (Borges et al., 2003; Harries and Bassey, 1990; Pontaga, 2004; Wagner et al., 1992).

We speculate that the shoulder’s torque-velocity profile appears more gradual than the lower body since the muscles spanning the hip and knee are larger and stronger to accommodate their functional demands, generating higher torques over a smaller range of velocities compared to the shoulder. Interestingly and contrary to our prediction, the torque-velocity relationships of the torso did not statistically differ between the constrained and unconstrained conditions. When exploring this further, we observed that there was a statistical difference between torso conditions for the fastest angular velocities across participants, where the mean across participants was 574 deg s ^-1^ in the constrained condition and 506 deg s ^-1^ in the unconstrained condition (p = 0.0015). Meanwhile, there was no statistical difference between conditions for the slowest angular velocities (p = 0.57), torques at the slowest angular velocities (p = 0.18), and torques at the fastest angular velocities (p = 0.06). The marginal difference for the torques at the fastest angular velocities with mean torques of 28 Nm and 30 Nm for constrained and unconstrained, respectively, indicate that the unconstrained condition may reduce the effort at the shoulder while presumably performing the task with similar linear velocity. In other words, the unconstrained condition may have allowed the participants to increase their torque output while decreasing the necessary angular velocity to achieve it. This would agree with past literature demonstrating the importance of pelvis and trunk rotation in increasing ball velocity for baseball pitching (Pappas et al., 1985; Stodden et al., 2001).

Due to the unique nature of our multiplanar task, drawing direct comparisons with existing angular velocity and torque values is challenging. The mean values of maximum angular velocity across all participants (574 deg s ^-1^ and 506 deg s ^-1^ for constrained and unconstrained, respectively) fall within the reported range of maximal horizontal adduction in prior studies (Escamilla et al., 1998; Feltner and Dapena, 1986; Matsuo et al., 2001; Naito et al., 2019; Seminati et al., 2015). Although the fastest angular velocities observed across participants (784 deg s ^-1^ and 671 deg s ^-1^ for constrained and unconstrained, respectively) are greater than that typically found for horizontal adduction, this is likely due to some contribution of internal rotation during the task which has been reported to achieve above 9000 deg s ^-1^ in professional baseball pitchers (Pappas et al., 1985). Previous research has documented shoulder torques between 30-50 Nm for internal rotation (Moghadam et al., 2011; Zanca et al., 2011), 16-54 Nm for abduction (Moghadam et al., 2011; Whitcomb et al., 1995), and 21-32 Nm for horizontal adduction (Moghadam et al., 2011). While these past studies are predominantly reported for isometric trials or single-plane motion, they offer a reference point for our multi-segment and multidimensional torques. A large proportion of our torque results at the slowest angular velocity (approximately 200 deg s ^-1^) are greater than the previously stated literature values. We speculate that our elevated torque data is a result of the movement itself allowing for multiple muscles to contract together, generating higher torques in contrast with studies that isolated specific motions and muscles (i.e. scaption to target the deltoid muscle). We also believe that extracting the torque and angular velocity at the instant of peak power may have allowed the muscles to be more favourably positioned. Selecting our metrics at the instant of peak power (which could vary across trials) is a distinct difference from the literature where the torque data was documented for a specific shoulder angle (ex. 90° shoulder abduction, 90° arm flexion).

### Power-velocity relationship

Shoulder power increased positively with angular velocity for the majority of participants in both torso conditions (Figure 5C, D). Interestingly, this reveals that the shoulder power in these participants remains on the ascending limb of the power-velocity curve, indicating that absolute maximal power may not have been reached. Prior studies have reported similar findings in the lower limb and hypothesized that velocities at which the absolute maximum power would occur were never reached during testing (Rahmani et al., 2001; Taylor et al., 1991). While we tested 1 m s ^-1^ above each participant’s maximum cable speed prior to the collection, perhaps the participants were able to adjust their upper limb joint contributions to increase power while diminishing the velocity demand specifically at the shoulder. Previous studies have proposed elastic energy storage and release at the shoulder during the arm-cocking phase (prior to arm acceleration) to generate the power necessary for high-speed throwing in humans (Roach et al., 2013). This mechanism likely did not impact the results of our investigation as the start position of the task did not involve pre-stretching/winding of the arm. Additionally, we saw no statistical difference between the power-velocity slopes from one condition to the other, though we found that the unconstrained condition resulted in a higher power-velocity slope, on average, and more participants (11) with statistically significant power-velocity relationships compared to the constrained (6). This may suggest that restricting torso rotation limits the ability of the shoulder to generate power, causing more variation in angular velocity and torque and therefore, linear regressions with flatter slopes. Varying compensatory mechanisms may have been at play to achieve the task while torso rotation was restricted. Examples include muscle activation patterns, joint posture, and reliance on muscle strength, all of which can vary among participants and affect the predictive power of the regression.

### Fatigue and learning effects

We conducted the maximum cable force and cable speed tests prior to and following the main data collection for each participant to assess if muscular fatigue or neuromuscular learning effects influenced the results. A paired t-test showed no statistical difference between the maximum force test that was conducted before and after the main data collection (p = 0.29) (Figure 7, left). This coupled with the statistical difference (p = 0.31e-5) between pre- and post-collection maximum cable speed (Figure 7, right) suggests that fatigue effects were unlikely at play. Since cable speeds were randomized, it is difficult to confirm if motor learning effects were present however, based on the maximum speed test, it is possible that motor learning of the repeated task and ample familiarization with the equipment may have aided in faster post-collection speeds. If this is true, it would support the importance of neuromuscular pathways and training to improve performance (Reiser, 2011).

### Limitations and future work

We recognize that the self-imposed torso restriction for the constrained condition could not allow for the complete isolation of the task to the shoulder joint. In the piloting phase of this study, we experimented with using a trunk brace during the task. While the brace limited trunk flexion in the sagittal and frontal planes, it did not reduce axial rotation. Previous studies have shown that the majority of the work required for high-velocity throwing is generated at the pelvis (Matsuo et al., 2001; Roach et al., 2013; Stodden et al., 2001). Given the equipment and nature of the task, it was impractical to perform the motion in a sitting position to eliminate the pelvis. As a compromise, the participants were instructed to keep their feet planted and pointed straight for the constrained condition. In addition, there exists inherent error in the kinematic calculations that originate from markerless motion capture and segment position estimation (Riemer et al., 2008). Future work should assess the accuracy of the kinematics resulting from optical motion capture with more accurate modalities such as biplanar video radiography (BVR). BVR allows the assessment of 3D, dynamic *in vivo* bone motion without soft tissue artifact (Bey et al., 2008; Kijima et al., 2015; Lawrence et al., 2021; Miranda et al., 2013; Tashman and Anderst, 2003). Using this novel motion capture system coupled with our outlined approach involving the instrumented cable machine would allow us to accurately measure specific joint contributions during rapid 3D shoulder motion. With this rich dataset, future work is also necessary to further investigate kinematic parameters such as how the instant of peak power and joint angles change throughout the range-of-motion depending on the test speed.

## Conclusions

We present a novel approach to examine shoulder mechanics and the intricacies of multiplanar upper limb movement through inverse dynamics and provide insights into the strength-speed trade-off. The results of this study allow us to more accurately describe the human shoulder’s biomechanical properties and capabilities, which are currently lacking in existing literature. While future work is necessary to analyze kinematic variables further and with higher accuracy, our torque-velocity and power-velocity analyses suggest that the shoulder may be able to redistribute loads and joint speeds during high-demand tasks to maintain linear speeds and reduce stress at the joint. This finding may aid our understanding of the upper limits the shoulder can withstand and the methods our neuromuscular system naturally seeks to maintain output while avoiding injury.

Although our shoulders may have evolved to handle high-speed throwing, early humans likely engaged in throwing activities far less frequently than today’s professional baseball pitchers, who typically throw 80-100 balls at maximum effort in a single game. Understanding the trade-offs between torque, power, and velocity can not only enhance sports performance through optimized training and inform injury prevention and rehabilitation, but also shed light on the evolutionary divergence of human shoulder morphology from non-human apes.

## Acknowledgements

We gratefully acknowledge our lab technicians, Kaito Hara-Lee and Dajung Yoon for their supportive, meticulous, and dedicated contributions during data collection and data processing. We also thank members of the Skeletal Observation Laboratory for their helpful feedback and insight. Lastly, we deeply appreciate the people who volunteered their time to participate in this study.

